# Whole genome methylation sequencing in blood from persons with mild cognitive impairment and dementia due to Alzheimer’s disease identifies cognitive status

**DOI:** 10.1101/2024.09.26.615196

**Authors:** Andy Madrid, Ligia A. Papale, Phillip E. Bergmann, Coleman Breen, Lindsay R. Clark, Sanjay Asthana, Sterling C. Johnson, Sündüz Keleş, Kirk J. Hogan, Reid S. Alisch

**Author notes:** Reid S. Alisch and Kirk J. Hogan contributed equally to this work. Co-author email addresses. **Correspondence:** Reid S. Alisch, Department of Neurological Surgery, University of Wisconsin-Madison School of Medicine and Public Health, 600 Highland Ave, Madison, WI 53792-3272, USA.; Kirk J. Hogan, Department of Anesthesiology, University of Wisconsin-Madison School of Medicine and Public Health, B6/319 Clinical Science Center, 600 Highland Ave., Madison, Wisconsin 53792-3272, USA.

## Abstract

**INTRODUCTION:** Whole genome methylation sequencing (WGMS) in blood identifies differential DNA methylation in persons with late-onset dementia due to Alzheimer’s disease (AD) but has not been tested in persons with mild cognitive impairment (MCI).

**METHODS:** We used WGMS to compare DNA methylation levels at 25,244,219 CpG loci in 382 blood samples from 99 persons with MCI, 109 with AD, and 174 who are cognitively unimpaired (CU).

**RESULTS:** WGMS identified 9,756 differentially methylated positions (DMPs) in persons with MCI, including 1,743 differentially methylated genes encoding proteins in biological pathways related to synapse organization, dendrite development, and ion transport. 447 DMPs exhibit progressively increasing or decreasing DNA methylation levels between CU, MCI, and AD that correspond to cognitive status.

**DISCUSSION:** WGMS identifies DMPs in known and newly detected genes in blood from persons with MCI and AD that support blood DNA methylation levels as candidate biomarkers of cognitive status.

## 1 BACKGROUND

Mild cognitive impairment (MCI) is an intermediate, prodromal diagnosis between cognitively unimpaired (CU) older adults and late-onset dementia due to Alzheimer’s disease (AD).[1] Persons with MCI are at elevated risk of progressing to dementia, with 80% converting to AD after 6 years.[1–7] While the molecular etiology of MCI and its transition to AD remains uncertain, recent evidence indicates that environmental influences increase the risk of AD in part through interactions with the epigenome.[8, 9] DNA methylation is an epigenetic modification that provides a gene-switching gate interposed between the unchanging genome and ever-changing environment essential to coordinate gene expression from conception to death. We recently reported that whole genome methylation sequencing (WGMS) in blood identifies thousands of differentially methylated positions (DMPs) in AD. The DMPs occur in 14% of coding genes (*N*=2,707), many with known biologically relevant roles in cognitive status and AD.[10] Accordingly, we integrate MCI in a model that compares whole genome DNA methylation levels in blood from persons with MCI with those from persons with AD and who are CU.

## 2 METHODS

### 2.1 Study participants

The experimental protocol was approved by the Institutional Review Board (IRB) of the University of Wisconsin School of Medicine and Public Health, Madison, WI. All participants signed an IRB-approved informed consent. Participants were enrolled in the Wisconsin Alzheimer’s Disease Research Center (WADRC) or the Wisconsin Registry for Alzheimer’s Prevention (WRAP) cohorts. Details of the study design and methods have been previously published.[10] Participants were clinically classified as cognitively unimpaired (CU) or as meeting criteria for MCI or AD based on National Institute on Aging-Alzheimer’s Association thresholds by consensus conference.[1, 11, 12] CU participants did not meet criteria for either MCI or AD. No participant provided a sample in more than one diagnostic category. Samples from persons whose clinical cognitive status reverted to CU, or who were diagnosed as having non-Alzheimer’s disease dementia at a subsequent visit, were excluded.

### 2.2 DNA extraction and generation of whole genome methylome data

Blood samples were acquired on the visit nearest after the date of MCI and AD diagnosis and matched for age by sample visit number in the CU cohort. A 10 mL whole blood sample was anticoagulated in EDTA and stored at −20^0^C. Genomic DNA was extracted using the Gentra Puregene Blood Core Kit C following the manufacturer’s protocol (Catalog# 158389, Qiagen^TM^, Hilden, Germany). Seven hundred nanogram samples of high molecular weight genomic DNA were forwarded to the University of Illinois at Urbana-Champaign Roy J. Carver Biotechnology Center for DNA sequence library construction using the NEBNext Enzymatic Methyl-seq (EM-seq™) kit (Ipswitch, MA)[13], and whole genome methylation sequencing on an Illumina NovaSeq6000 sequencer (San Diego, CA). Image processing and sequence extraction used the Illumina Pipeline (Illumina, San Diego, CA). Raw fastq sequence files were forwarded to Dr. Alisch’s laboratory via a secure website.

#### 2.3.1 Data processing

Data processing was performed with the ENCODE gemBS-based processing pipeline.[14] Raw WGMS data were analyzed with FastQC to ensure that all samples met quality control thresholds for sequencing reads and with Trim Galore to remove adapter content and trim sequence reads. DNA sequencing reads were aligned to the GRCh38.14 (hg38) human reference sequence[15] with gemBS in pair-ended mode with default settings. Methylated and unmethylated sequence read counts were merged across the 2 DNA strands of each sample. CpG loci were filtered to have median coverage of 5 or greater, with 50% or fewer missing values among all participants. CpGs on sex chromosomes were removed from analysis. A total of 25,244,219 CpGs were retained after filtering and entered into subsequent analyses for comparisons between MCI *vs*. CU, AD *vs.* MCI, and AD *vs*. CU. Missing values were imputed using diagnostic cohort averages at a specific CpG locus after merging across strands.

### 2.4 Statistical analysis

#### 2.4.1 Cell-type deconvolution

Whole blood comprises a composite of different white blood cell types each with a distinct methylation profile. To adjust for leukocyte heterogeneity, we used reference-free deconvolution of subpopulations existing in methylation data (DXM) to estimate the constituent fraction of each white blood cell type as previously reported.[16]

#### 2.4.2 Differential methylation analysis

Principal components (PC) were calculated on the 5% of CpGs with the greatest DNA methylation variability on each chromosome to detect potential batch effects between the WADRC and WRAP cohorts, age, and sex. Dispersion shrinkage for sequencing (DSS)[17] was used to fit a beta-binomial model for each CpG locus adjusted for estimated white blood cell proportions, the first 2 PCs, age, sex, and body mass index (BMI). All 382 samples were used for model fitting. A diagnosis-specific effect on methylation level for each CpG was tested with a Wald test of the corresponding regression coefficient for pairwise comparisons of 3 diagnoses of interest *i.e.,* MCI *vs*. CU, AD *vs.* MCI, and AD *vs*. CU. To identify linear trends in methylation levels, diagnostic identifiers were assigned numerical values during model fitting with CU treated as 0, MCI treated as 1, and AD treated as 2. The empirical distribution of the 25,244,219 *P*-values resulting from the theoretical null (standard normal distribution) for Wald tests from each comparison deviated from the expected uniform distribution with enrichment around small *P*-values. Accordingly, under-dispersed theoretical null test statistics were used to model the empirical null, and to calculate local false discovery rates (lFDRs) to correct for multiple testing with fdrtool.[18] For 2-way diagnosis comparisons (*i.e.,* MCI *vs*. CU, AD *vs.* MCI, and AD *vs*. CU) a CpG locus with a lFDR < 0.05 and an estimated methylation difference ≥2.5% was used to identify differentially methylated positions (DMPs). For 3-way cognitive status comparisons for analysis of trends from CU to MCI to AD, a CpG locus with a lFDR < 0.05 and an estimated methylation difference ≥2.5% between CU and MCI, between MCI and AD, and between CU and AD was identified as a DMP.

#### 2.4.3 Gene annotation, gene ontology analysis, and gene structure identification

5′ to 3′ genomic coordinates for all protein coding genes were obtained from ENSEMBL (v86).[19] Gene coordinates were extended to include 3 kilobases (kb) 5′ of a transcription start site (TSS) to 200 base pairs (bp) 3′ of a transcription termination site (TTS). Regions within 5 kb 5′ of a TSS and 200 bp 3′ of a TSS were utilized as gene promoters. Two criteria were used to identify differential methylation in genes and promoters. First, each gene or promoter must comprise a gene-wide or promoter-wide lFDR < 0.01. To calculate gene-wide or promoter-wide lFDRs, *P-*values of each CpG within the gene or promoter were aggregated into a harmonic mean *P*-value.[20] The harmonic mean *P*-value controls the family-wise error rate of multiple statistical tests and does not assume independence among tests. The harmonic mean *P*-value thereby provides a preferred framework for aggregating *P*-values to correct for gene size (*i.e*., number of CpGs/gene), and for correlated methylation levels of neighboring CpGs. This step was repeated for each gene and promoter yielding a single *P*-value for the whole gene or promoter. lFDRs were then calculated from the *P*-values to provide gene-wide and promoter-wide false discovery rates (FDR) that account for multiple testing. The second criterion to test for differential methylation of genes and promoters required at least one DMP (defined above) to be within the gene or promoter. Accordingly, genes and promoters with an lFDR < 0.01 and at least 1 DMP are identified as differentially methylated genes and promoters. Of note, annotation may assign a DMP to more than a single gene. Gene Ontology (GO) analyses were performed with clusterProfiler (version 4.6.0)[21] that conducts an over/under-representation test.[20, 22]

#### 2.4.4 Differential methylation analysis of blood promoter–enhancer interactions

To identify differential methylation in gene promoter and enhancer regions, promoter–enhancer interactions in white blood cells were examined using published Promoter Capture Hi-C (PCHi-C) data.[23] An interaction was deemed significant if the minimum CHiCAGO score across all blood cell types was > 5. Interactions were filtered to have a median CHiCAGO score > 5 across all blood cell types to assure adequate signal. After removal of false positive (*e.g*., bait–bait) and sex chromosome interactions, remaining promoter/enhancer interactions were lifted over to the human genome reference version GRCh38 with the UCSC liftOver tool.[24] This approach retained 104,554 promoter–enhancer interactions comprising 9,631 distinct promoters and 58,654 distinct enhancers for comparison to WGMS data. These promoters and enhancers were considered differentially methylated if they contained one or more DMPs. Over- or under-representation of DMPs in enhancer regions were tested against a null distribution generated by random sampling of a set of the same number of DMPs from the 25,244,219 CpGs. The empirical null distribution quantified the proportion of DMPs located within a blood enhancer based on 10,000 random Monte Carlo samplings. A 2-tailed test statistic was calculated to ascertain the number of DMPs occurring within an enhancer by chance alone. The same simulation procedure was also used to identify under- or over-representation of DMPs in promoters.

#### 2.4.5 Comparisons of differential DNA methylation with differential expression in AD

To evaluate whether genes defined as above with 1 or more DMPs in 1 or more promoter– enhancer interactions were differentially expressed between CU, MCI and AD, WGMS data were compared to published transcriptome data that identified 846 differentially expressed genes in blood from persons with AD.[25] After removal of differentially expressed genes on sex chromosomes, 790 were retained for subsequent analyses. Differentially expressed genes were then correlated with differentially methylated genes, differentially methylated enhancer genes, and differentially methylated promoter genes by gene symbol.

#### 2.4.6 DNA methylation level trends across cohort cognitive status

After differential methylation analysis by DSS, adjusted DNA methylation levels per sample were tested for trend by the Cuzick trend test.[26] Cognitive status groups were permuted 100 times to obtain *P*-values, followed by correction for multiple testing by Benjamini-Hochberg FDR. An FDR threshold of <0.05 from Cuzick *P*-values, coupled with ≥2.5% methylation difference between diagnostic groups and an lFDR threshold of 0.05 from *P*-values from DSS were identified as DMPs.

#### 2.4.7 Software for statistical analysis and data visualization

All statistical analyses were conducted in R (v4.2.0). Figures were generated with ggplot2 (v3.4.2)[28], ggsci (v3.0.0)[29], and cowplot (1.1.1).[30]

## 3 RESULTS

### 3.1 Study participants

WGMS was performed on 99 WADRC and WRAP participants with MCI for comparison with CU (174) and AD (109) participants.[10] Fifty-two percent of MCI and CU participants, and 43% of AD participants, were female (Table 1). MCI participants were approximately 4 years older than CU participants with approximately 1.5 fewer years of education and no significant difference in body mass index. All but 11 MCI participants reported European ancestry. Forty-eight MCI participants were apolipoprotein E (*APOE*) ε4 heterozygotes and 12 were *APOE* ε4 homozygotes.

**Table 1:**
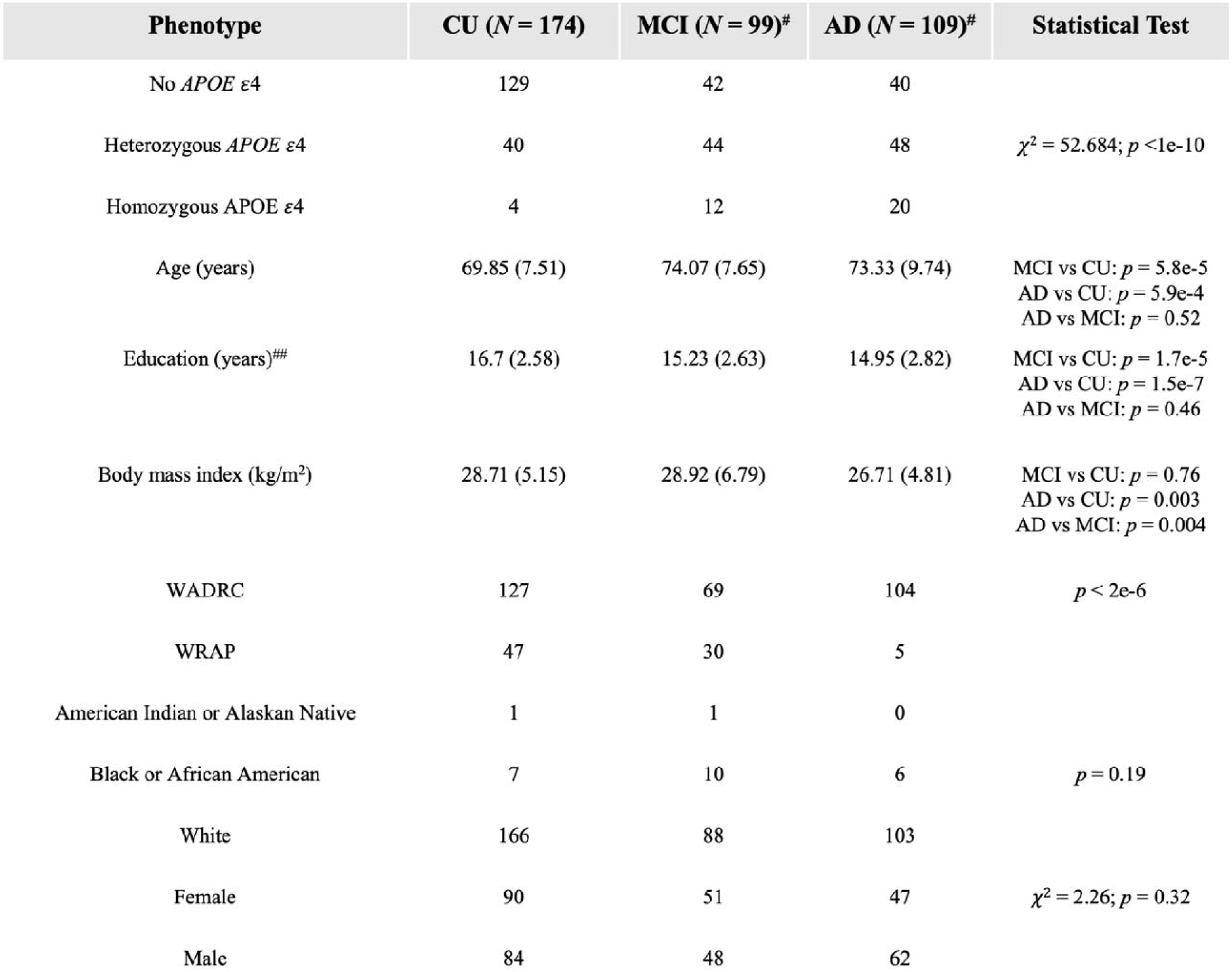
Means and (standard deviations) for persons who are cognitively unimpaired (CU), and persons with mild cognitive impairment (MCI) and late-onset dementia due to Alzheimer’s disease (AD) demographic variables. *t*-statistics and corresponding *P-*values without correction for multiple testing are reported for pairwise comparisons. A Chi-squared test statistic was used to test independence for *APOE* heterozygote and homozygote status. Three participants (one from each group) were not genotyped and are not included in statistical test for *APOE* status (^#^). Two MCI participants were missing education data (*i.e*., all years of formal education) and were not included in the statistical test for Education (^##^). Fisher’s exact test was used to test independence for self-identified ancestry with zero counts in some groups. Abbreviations: *APOE*: apolipoprotein E; CU: cognitively unimpaired; MCI: mild cognitive impairment; AD: late-onset dementia due to Alzheimer’s disease.

### 3.2 Whole genome DNA methylation detection

After filtering sequenced reads for quality assurance, EM-seq generated an average of 561 million paired-end reads for each participant (CU=174, MCI=99, and AD=109), and an average of 539 million sequence reads uniquely mapped to the human genome (GRCh38.14 [hg38]). The unique reads provide an average genomic coverage of 53x, with a mean of 38x after merging across strands (Figure S1). Inclusion of MCI data for the 3 pairwise comparisons added CpGs below the filtering criteria thresholds (*i.e*., sites with > 50% missing data or median coverage less than 5x) resulting in an overall lower number of CpGs that meet quality assurance metrics than in the single pairwise comparison of AD *vs*. CU described in an earlier report.[10] Nonetheless, these data provided interrogation of 25,244,219 CpGs in a 3 pairwise comparison model. Batch effects in WGMS data were tested with the first 2 principal components calculated on the 5% of CpGs with the greatest DNA methylation variability on each chromosome (*e.g*., PC1 and PC2 carried 5.1% and 2.5%, respectively, of the total variability on chromosome 1). No batch effects for participant cohort (WADRC or WRAP), cognitive diagnosis, sex, or reported ancestry were observed (Figure S2). These data were used for the following 2-way diagnosis comparisons: MCI *vs*. CU; AD *vs.* MCI; and AD *vs*. CU to identify differentially methylated positions (DMPs).

### 3.3 Differential DNA methylation between CU and MCI

Differential DNA methylation in blood from CU and MCI participants were observed at 9,756 CpG loci (Figure 1A; Table S1). Over two thirds (*N*=6,536) of the MCI-associated DMPs are hypermethylated (*i.e*., higher DNA methylation levels in MCI than CU participants). MCI-associated DMPs are distributed across all recognized genomic structures, including 49.8% within a gene intron and 15.2% within 5 kb of transcription start sites (TSS, Figure 1B). These data suggest that the human genome is hypermethylated in persons with MCI.

**Figure 1:**
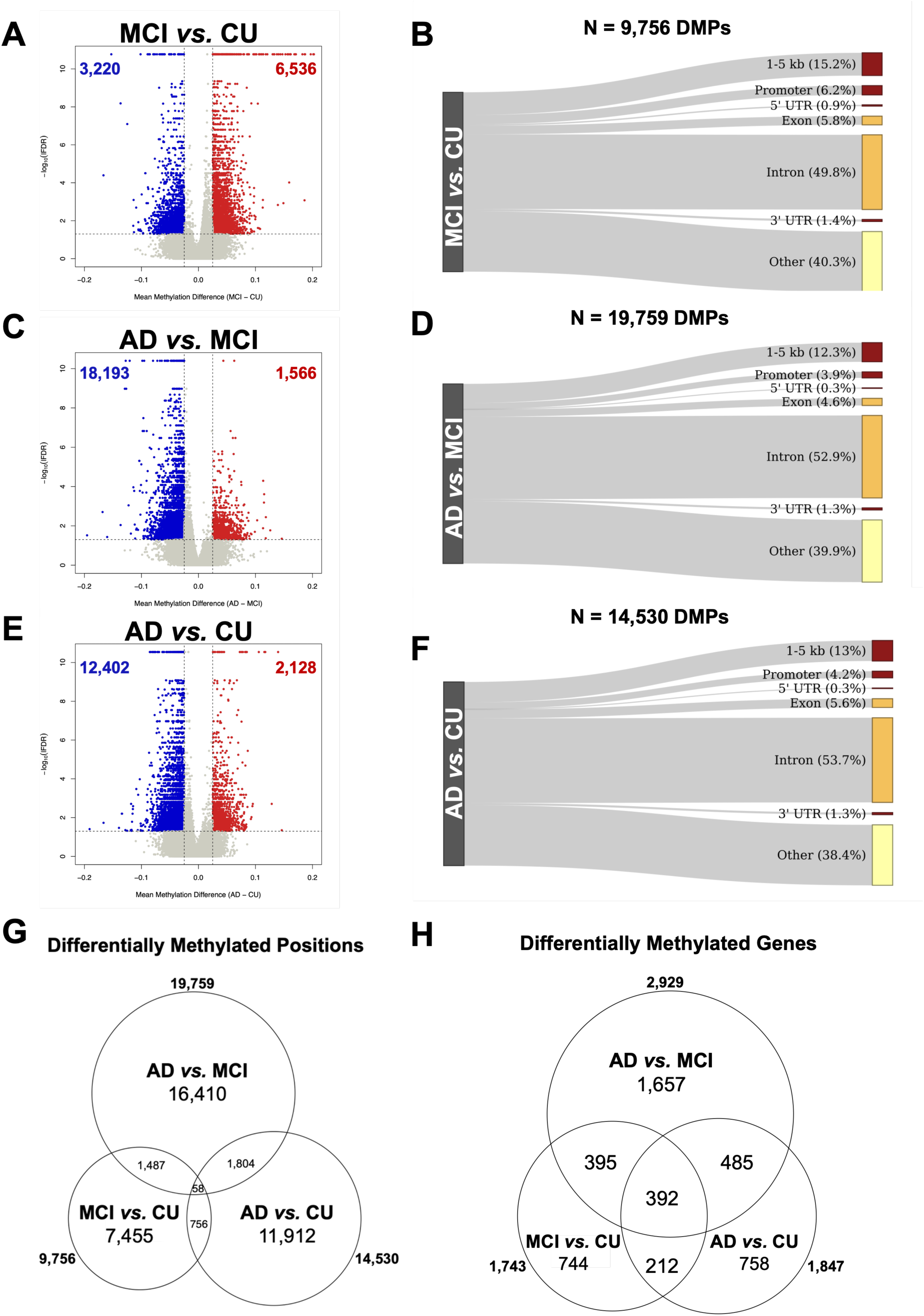
Identification of 5’-cytosoine-phosphate-guanine-3’ (CpG) loci that comprise differentially methylated positions (DMPs) in blood between participants who are cognitively unimpaired (CU) and those with mild cognitive impairment (MCI), between those with MCI and those with late onset dementia due to Alzheimer’s disease (AD), and between those with AD and those who are CU. Volcano plots of 25,244,219 WGMS CpG loci show the difference in mean methylation percent (x-axis) and the significance (y-axis, local false discovery rate (lFDR) between persons with CU and MCI (A), between persons with MCI and AD (C), and between persons with CU and AD (E). Each point corresponds to a single CpG locus: not differentially methylated(grey); hypomethylated (blue); and hypermethylated (red). Strings of blue and red points across the top of the plots have reached the minimum lFDR as computed by fdrtool.[18] Sankey plots show the locations of 9,756 MCI *vs.* CU DMPs (B) 19,759 AD *vs.* MCI DMPs (D), and 14,530 AD *vs.* CU DMPs (F) DMPs relative to genomic structures comprising the following: 1-5 kilobases (kb) upstream of the transcription start site (TSS; 1-5 kb); up to 1kb upstream of the TSS (promoter); the 5’ untranslated region (5’ UTR); within a gene exon (exon); within a gene intron (intron); the 3’ untranslated region (3’ UTR); and 5kb or more from any gene structure (Other). 3-way Venn diagrams showing the overlaps of differentially methylated positions (G) and genes (H) between MCI *vs*. CU and (9,756 and 1,743, respectively), AD *vs*. MCI (19,759 and 2,929, respectively), and AD *vs*. CU (14,530 and 1,847, respectively). Fifty-eight DMPs and 392 differentially methylated genes are shared between the 3 pairwise comparisons.

### 3.4 Differential DNA methylation between MCI and AD

Differential DNA methylation in blood from MCI and AD participants were observed at 19,759 DMPs (Figure 1C; Table S1). The majority of DMPs between MCI and AD are hypomethylated (92%, *i.e*., lower DNA methylation levels in AD than MCI participants). DMPs between MCI and AD are distributed across all recognized genomic structures, including 52.9% within a gene intron (Figure 1D). When these DMPs were compared to AD genomic risk loci reported in a large genome-wide association study (GWAS)[31], 36% (27/75) of the AD genomic risk loci comprised at least one DMP between MCI and AD (Table S2), suggesting DMPs are located in known risk loci of AD. These data indicate that the molecular differences between MCI and AD are greater than between CU and MCI.

### 3.5 Differential DNA methylation between CU and AD

Differential DNA methylation in blood from CU and AD participants were observed at 14,530 DMPs (Figure 1E; Table S1). The number of AD-associated DMPs in the present data differs from that previously reported[10] because the addition of 99 samples from participants with MCI generated improved statistical model fitting and the DMP classification employed a greater stringency (*i.e*., a ≥2.5% change in DNA methylation level). Nonetheless, as previously reported, the majority of AD-associated DMPs are hypomethylated (85%, *i.e*., lower DNA methylation levels in AD than CU participants). These DMPs are distributed across all recognized genomic structures, including 53.7% within a gene intron (Figure 1F). Thirty-seven percent (28/75) of reported AD genomic risk loci[31] comprise at least one DMP between CU and AD (Table S2).

### 3.6 Differentially methylated positions (DMPs) shared between MCI and AD

Fifty-eight DMPs are shared between the 3 comparisons (*i.e.,* MCI *vs*. CU, AD *vs.* MCI, and AD *vs*. CU, Figure 1G). Fifty are hypermethylated and 8 are hypomethylated in MCI and AD participants compared to CU (Table S1) and the majority (90%) have higher DNA methylation levels in MCI than in AD participants. Four of the 58 DMPs annotate to genes, including 3 DMPs in thymine DNA glycosylase (*TDG*) that are hypomethylated in both MCI and AD, and a single DMP in the gene encoding fragile histidine triad diadenosine triphosphatase (*FHIT*). The remaining 54 shared DMPs are located distally from any protein coding gene, with 44/54 located in recognized repetitive DNA sequences.

### 3.7 Differentially methylated genes between MCI and AD

Genomic annotation of differential DNA methylation associated with MCI and AD identifies unique and shared genes from all 3 pairwise comparisons (*i.e*., MCI *vs.* CU, *N*=1,743; AD *vs.* MCI, *N*=2,929; and AD *vs.* CU, *N*= 1,847; Figure 1H; Table S3). To test whether DNA methylation levels in blood differ in genes associated with the onset and progression of early- and late-onset Alzheimer’s disease, we examined differential DNA methylation in well-known AD genetic susceptibility genes including Apolipoprotein E (*APOE*)[32], amyloid beta precursor protein (*APP*)[33], microtubule associated protein tau (*MAPT*)[34], presenilin 1 (*PSEN1*)[33], presenilin 2 (*PSEN2*)[33], and translocase of outer mitochondrial membrane 40 (*TOMM40*)[35] (Figure 2). None of the AD-associated genes are differentially methylated between CU and MCI. In contrast, *APP*, *MAPT*, and *TOMM40* are differentially methylated between MCI and AD, comprising 5, 1, 2 DMPs, respectively (Figure 2). *MAPT* and *TOMM40* are differentially methylated between CU and AD, comprising 9 and 1 DMPs, respectively (Figure 2). *PSEN2* contains a single DMP between CU and AD but did not meet criteria for a differentially methylated gene (see section 2.4.3). *APOE* and *PSEN1* are not differentially methylated in any comparison.

**Figure 2:**
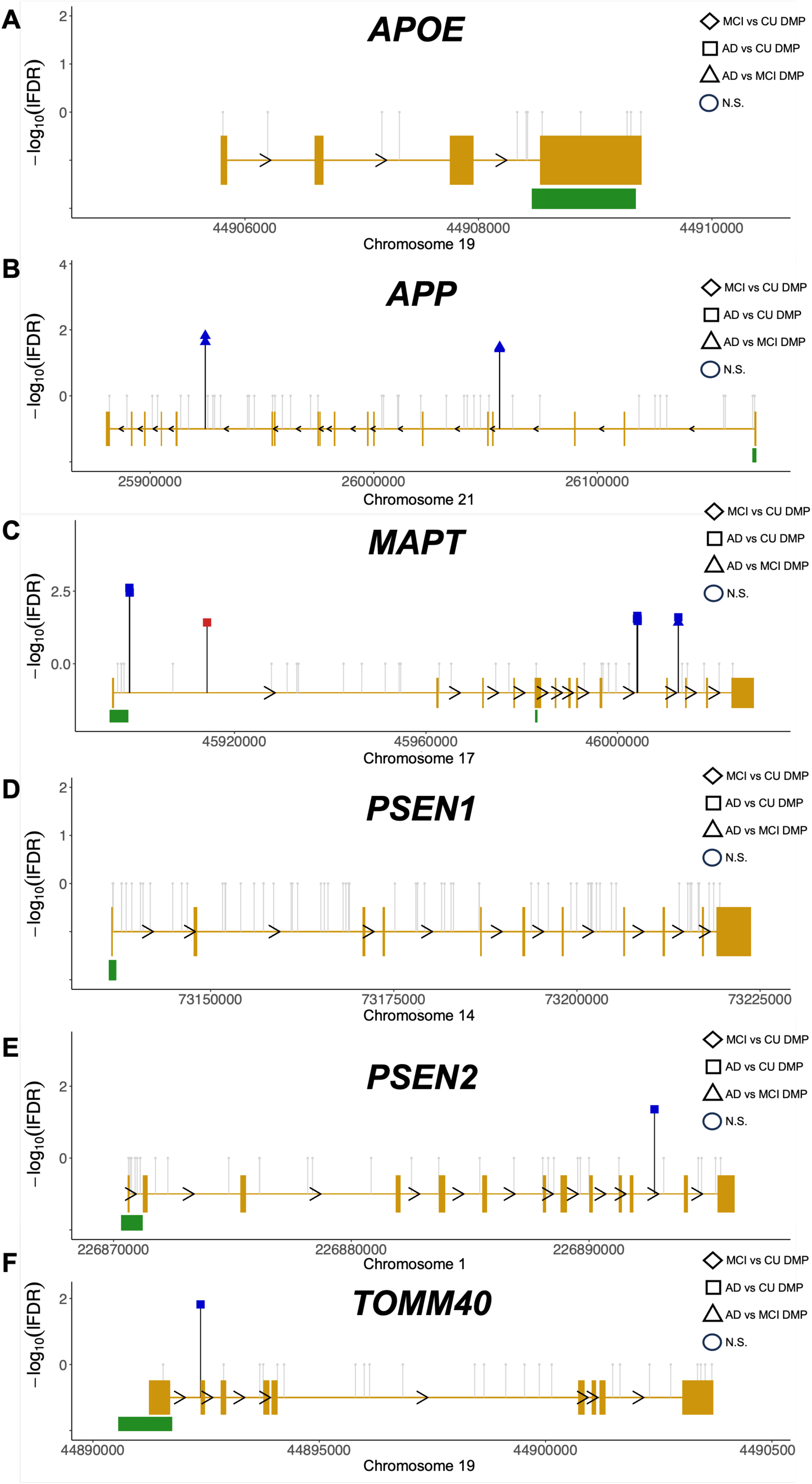
Genes with 5′-cytosoine-phosphate-guanine-3′ (CpGs) that are differentially methylated positions (DMPs) in recognized Alzheimer disease-related genes. Schematic diagrams of the methylome landscapes of representative AD-associated genes (A) – (F) Sense (>) and anti-sense (<) strands are depicted with gene name abbreviations above each panel (see section 2.4.3 for gene nomenclature). Alignment to human reference genome hg38 coordinates are displayed in basepairs (bp) at the bottom of each panel with the chromosome number (x-axis). The direction of gene transcription and introns are depicted by small black arrowheads on the gold line. Gold rectangles indicate coding exons connected by a gold line that indicate introns. CpG islands > 300bp in length are indicated by green rectangles. The relative location of CpG dinucleotides in each gene is shown as a vertical line with a geometric shape on top. Significance (y-axis) of differentially hypermethylated CpGs (red shapes) and hypomethylated CpGs (blue shapes) are shown for MCI *vs*. CU (diamond), AD *vs*. MCI (triangle), and AD *vs*. CU (square). One in 25 non-significant CpGs in each gene are shown (grey vertical line with circle, N.S.). A corrected significance level of lFDR < 0.05 and >2.5% differential methylation level was adopted for all comparisons.

Multiple genes are differentially methylated in all 3 comparisons, many with previously reported links to AD and neurodegeneration including glutathione S-transferase mu 1 (*GSTM1*)[36], RNA-binding fox-1 homolog 1 (*RBFOX1*)[37], and thymine DNA glycosylase (*TDG*)[38] (Figure 3). *GSTM1*, *RBFOX1*, and *TDG* comprise 15, 19, and 15 DMPs between CU and MCI, respectively; 10, 29, and 4 DMPs between MCI and AD, respectively; and 13, 17, and 11 DMPs between CU and AD, respectively.

**Figure 3:**
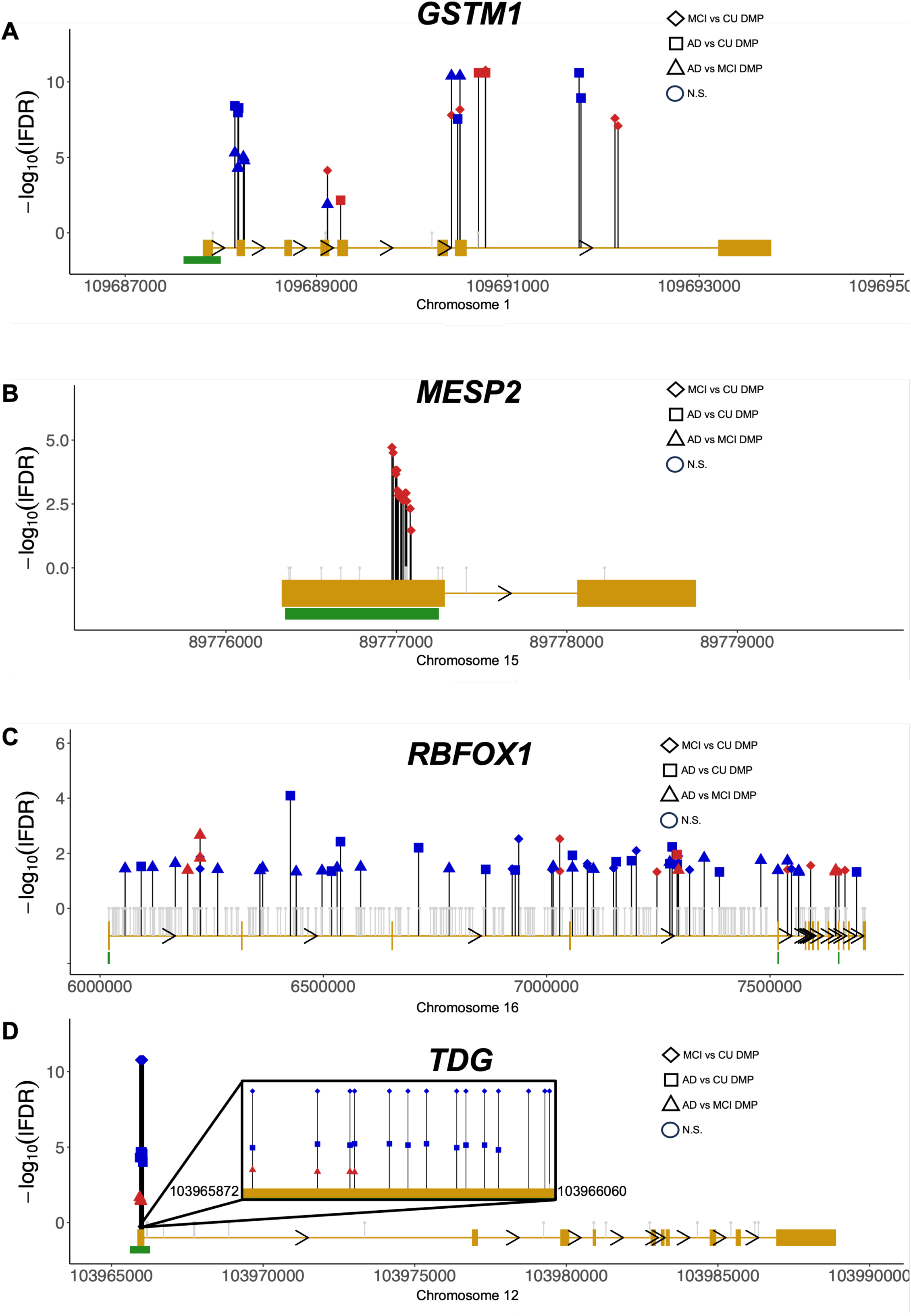
Genes with DMPs between persons with and without MCI and AD. Schematic diagrams of the methylome landscapes of representative AD-associated genes (A) – (D) Sense (>) and anti-sense (<) strands are depicted with gene name abbreviations above each panel (please see section 2.4.3 for gene nomenclature). Alignment to human reference genome hg38 coordinates are displayed in basepairs (bp) at the bottom of each panel with the chromosome number (x-axis). The direction of gene transcription and introns are depicted by small black arrowheads on the gold line. Gold rectangles indicate coding exons connected by a gold line that indicate introns. CpG islands > 300bp in length are indicated by green rectangles. The relative location of CpG dinucleotides in each gene is shown as a vertical line with a geometric shape on top. The significance level (y-axis) of differentially hypermethylated CpGs (red shapes) and hypomethylated CpGs (blue shapes) are shown for MCI *vs*. CU (diamond), AD *vs*. MCI (triangle), and AD *vs*. CU (square). One in 25 non-significant CpGs in each gene are shown (grey vertical line with circle, N.S.). A corrected significance level of lFDR < 0.05 and >2.5% differential methylation level was adopted for all comparisons. The inset window in panel D depicts the DMPs located in the *TDG* promoter in higher resolution, including the flanking genomic coordinates of the *TDG* promoter.

### 3.8 Differentially methylated gene ontology pathways in MCI and AD

A gene ontological analysis was computed on the hypermethylated (*N*=1,260) and hypomethylated (*N*=693) genes to examine the biological relevance of the genes that are differentially methylated between persons with CU and MCI. Multiple gene ontology terms with biological relevance to cognition were identified, including axon development, synapse assembly, dendrite development, and potassium ion transport (Figure 4A, Table S6). Numerous differentially methylated genes identified in these pathways have previously recognized links to AD (*e.g*., *ANK3* and *HLA-DQA1*[31]), indicating that common genes and pathways participate in the pathogenesis of MCI and AD. These findings were recapitulated in the gene ontological analysis of hypermethylated (*N*=358) and hypomethylated (*N*=2,777) genes found between persons with MCI and AD. Biologically relevant terms that are shared between CU and MCI and between MCI and AD include synapse assembly and potassium ion transport (Figure 4A).

**Figure 4:**
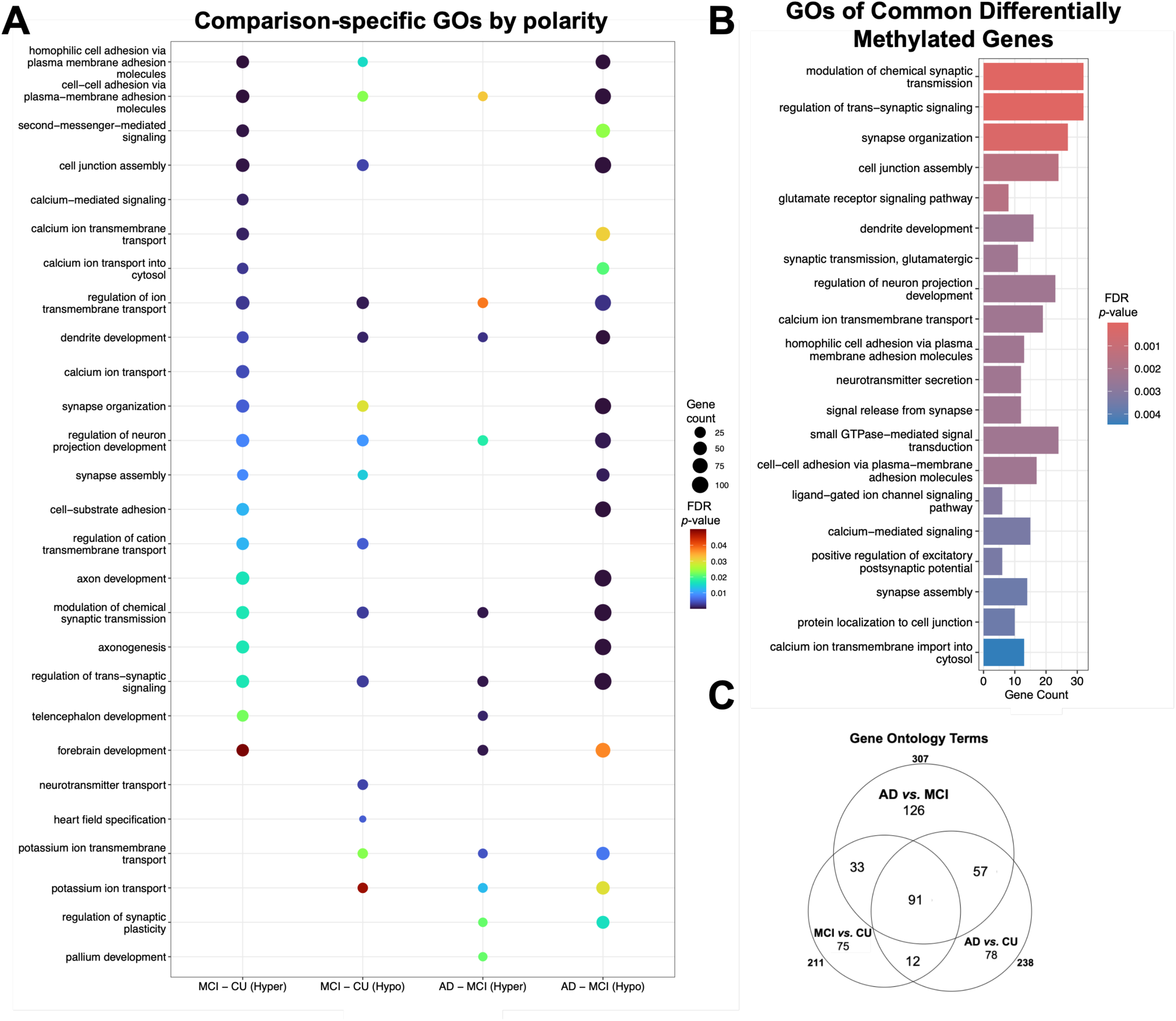
Gene ontological enrichment analyses and overlap. (A) Comparison-specific gene ontologies (GO) by diagnosis and by hypermethylation *vs*. hypomethylation. The GO plot shows genes comprising one or more DMPs between persons with CU and MCI, and between persons with MCI and AD. The top terms by significance (y-axis by false discovery rate [FDR] *P*-value < 0.05) are depicted by the number of differentially hypermethylated (Hyper) and hypomethylated (Hypo) genes in each term as provided by the diameter of each disc and for each comparison (x-axis). (B) GOs of common differentially methylated gene. The top 20 terms by significance are depicted (y-axis by local false discovery rate [lFDR] < 0.05). The number of contributory genes (x-axis) associated with each term is provided together with the color-coordinated FDR *P*-value. (C) Overlap of gene ontology terms. The 3-way Venn diagram illustrates the overlap of GO terms between MCI *vs*. CU (211), AD *vs*. MCI (307), and AD *vs*. CU (238). Ninety-one GO terms are shared between the 3 pairwise comparisons.

A gene ontological analysis of the shared genes from all 3 pairwise comparisons (Figure 1H, *N*=392 genes) retrieved additional terms with biological relevance to cognition through synaptic plasticity and glutamate signaling (Figure 4B), further underscoring shared genes and pathways. Thirty percent to 43% of GO terms are shared between the 3 pairwise comparisons (Figure 4C). These data indicate that multiple molecular processes are shared between persons with MCI and AD despite distinct clinical phenotypes.

### 3.9 Differential DNA methylation in gene promoters and enhancers between CU and MCI and between MCI and AD

To test whether DMPs located more than 5kb beyond a gene boundary and its putative promoter (*i.e*., >40% of all DMPs) contribute to altered gene expression, we interrogated 58,654 published blood-specific enhancer-promoter interactions[39] for enrichment of DMPs between CU and MCI, and between MCI and AD. We identified 387 differentially methylated enhancers and 212 differentially methylated promoters with one or more DMPs between CU and MCI (Table S4), and 666 differentially methylated enhancers and 315 differentially methylated promoters between MCI and AD (Table S5). Comparison of these data with published differential RNA expression data in blood from persons with AD[25] shows that 59 of the 666 differentially methylated enhancers between MCI and AD have long-range interactions with 53 promoters of genes known to be differentially expressed in blood of AD participants, including thioredoxin-interacting protein (*TXNIP*, Figure 5)[40]. These data suggest DMPs located beyond gene boundaries contribute to altered gene expression.

**Figure 5:**
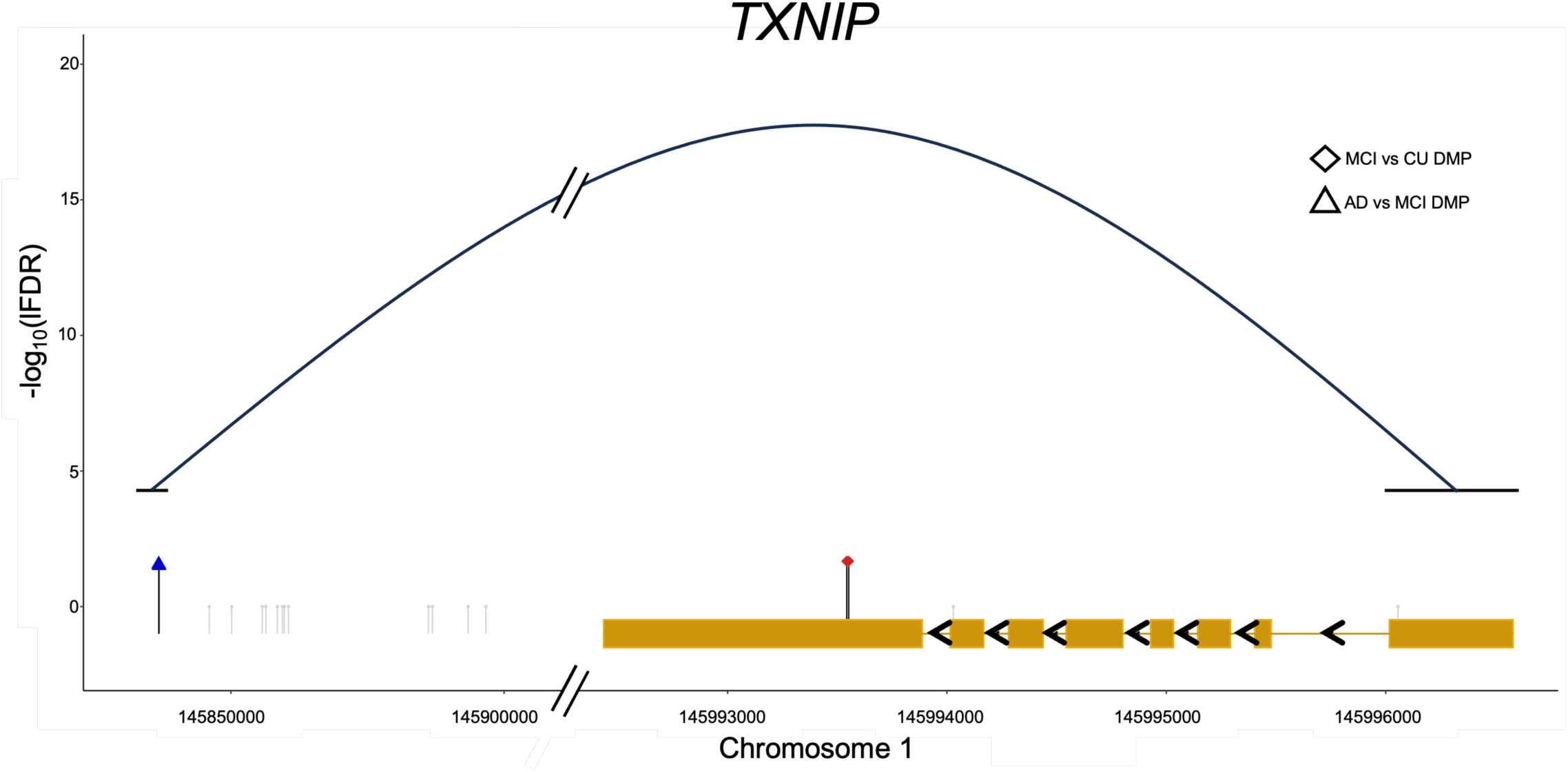
The long-range interaction between a differentially methylated enhancer and a gene promoter that is differentially expressed in the blood of persons with AD.[25] The antisense strand is shown for *TXNIP* (Thioredoxin-Interacting Protein). Alignments to human reference hg38 genome coordinates are displayed in bp at the bottom of the panel along chromosome 1. The direction of gene transcription and introns are depicted by small black arrowheads on the gold line. Gold rectangles indicate coding exons linked by a gold line that indicate introns. The relative location of 2 CpG dinucleotides in are shown as a vertical line with a geometric shape on top. The significance levels (y-axis) of a differentially hypermethylated CpG (red shape) and a hypomethylated CpG (blue shape) are shown for MCI *vs*. CU (diamond) and AD *vs*. MCI (triangle). One in 25 non-significant CpGs are shown (grey vertical line). A corrected significance level of lFDR < 0.05 was adopted for all comparisons. The arc indicates the enhancer region that interacts with the *TXNIP* promoter.

### 3.10 Progressive increases and decreases in DNA methylation levels of shared DMPs

To test whether DNA methylation levels change progressively at specific DMPs in tandem with changes in cognitive status, we tested for DMPs with increasing or decreasing DNA methylation levels across the 3 cohorts and found 447 DMPs with significant increments (*N*=125, Figure 6A) or decrements (*N*=322, Figure 6B) in DNA methylation levels (FDR < 0.05). The 447 DMPs comprised 87 DMPs with decreasing DNA methylation levels and were annotated to 99 genes (Table S3). These data indicate that blood DNA methylation levels may anticipate changes in cognitive status due to AD.

**Figure 6:**
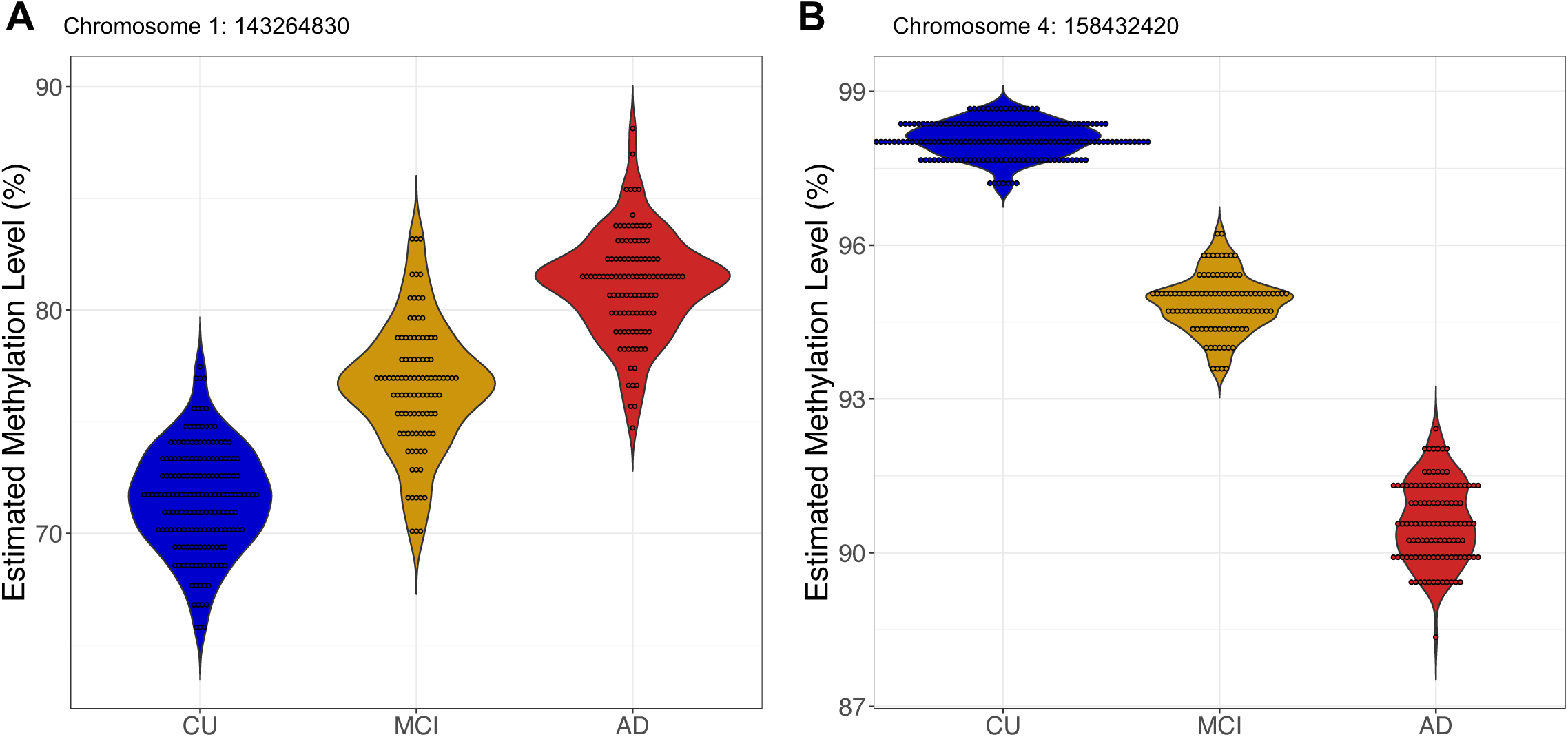
DMPs with increasing and decreasing methylation in separate cohorts between CU, MCI and AD. (A-B) Violin plots show the diagnosis (x-axis) and the percent methylation level (y-axis) for DMPs with significant increments (A) and decrements (B) in DNA methylation levels between persons with CU, MCI and AD. Dots within the violin plots represent the percent methylation level of each participant in each cohort.

## 4 DISCUSSION

We recently reported that whole genome methylation sequencing (WGMS) in blood identifies many thousands of CpG sites that are differentially methylated in persons with AD.[10] Reports of DNA methylation levels in blood samples from persons with MCI have been limited by the use of the Illumina HumanMethylationEPIC array, which interrogates just 4% of the human genome.[41] Here we tested whether persons with MCI have differential DNA methylation compared to persons with AD and those who are CU using WGMS to interrogate all the CpG sites in the human genome. We identified 9,756 DMPs comprising 0.04% of 25,244,219 candidate CpG sites for differential methylation in the human genome after adjusting for demographic variables (Table 1) and correction for multiple testing (Table S1). In contrast to global hypomethylation and regional hypermethylation of the human methylome that has been reported with aging,[42] greater than 66% of MCI-associated DMPs are hypermethylated, suggesting that MCI-associated DMPs correspond to changes that are not generic to aging unlike the AD-associated DMPs (Figure 1).[43] This shift in DNA methylation abundance from MCI to AD is confirmed by our finding that 92% of DMPs between MCI and AD are hypomethylated (Figure 1).

Genomic annotation of differential DNA methylation in blood from persons with MCI identifies 1,743 differentially methylated genes comprising nearly 10% of genes in the human genome.[44] Although our data are derived from blood samples, numerous identified DMPs are in genes that are known to play key roles in neuronal function. Other reports have identified a remarkable concordance between DNA methylation patterns in circulating white blood cells and the brain,[41, 45–49] in keeping with the relevance of our findings to phenotypes expressed in the central nervous system. Gene ontological analyses of the 1,743 genes that are differentially methylated in MCI show that the great majority of terms exhibit biological relevance to cognitive status, including terms linked with synaptic plasticity and regulation of neurotransmission (*e.g.,* glutamatergic neurotransmission). Gene ontological analysis of the differentially methylated genes that are shared between the 3 pairwise comparisons (*N*=392, Figure 1H) reports highly enriched terms with biological relevance to MCI and AD indicating that common pathways participate in the pathogenesis of both MCI and AD (Figure 5C). Multiple pathways among the 392 shared genes contribute to cognitive status including 21 synapse mechanism terms and 5 glutamatergic transmission terms. These data indicate that despite a clinically distinct phenotype between persons with MCI and AD, shared blood-based DNA methylation biomarkers of AD are present in MCI individuals.

Differentially methylated genes observed in the 3 pairwise comparisons participate in pathways with recognized links to AD including variants in gamma-aminobutyric acid type A (GABA) synthesis (*i.e*., *GABRA3*) and the glutamate ionotropic receptor NMDA family of genes (*e.g*., *GRIN2A*, *GRIN2B*, and *GRIN3A*) that encode for N-methyl-D-aspartate (NMDA) glutamatergic receptor units.[50] Variants of these genes alter the synaptosomes of brains from persons with AD, in keeping with an imbalance between excitatory and inhibitory synaptic functions. The RNA-binding protein RBFOX1 stabilizes mRNAs that encode synaptic transmission proteins that are degraded in postmortem brains of individuals with AD.[37] Sortilin-related VPS10 domain containing receptor 3 (*SORCS3*) is differentially methylated in all 3 pairwise comparisons, is differentially expressed in postmortem brain of persons with AD, and takes part in APP processing with critical roles in memory formation and synaptic plasticity.[51] A majority of shared differentially methylated genes have prior established links to MCI or AD (Table S3). Of note, multiple protein coding genes comprising DMPs that lack presently recognized links to AD pathogenesis directly participate in DNA methylation (*e.g*., *DNMT1*) and DNA de-methylation pathways (*e.g*., *TET2,* and *TDG*).[52]

Enhancers are *cis*- and *trans*-acting regions of the human genome that coordinate cell-type-specific gene expression by looping long distances to reach physical proximity with the promoters of their target genes.[39] Differential DNA methylation in enhancers corresponds to disease status in a diversity of clinical settings.[53–55] Here we identified 666 differentially methylated blood-specific enhancers[23] comprising 59 that interact with the promoters of 53 genes that are differentially expressed in blood samples from participants with AD.[25] These data suggest that differential DNA methylation also influence long-range gene expression related to MCI. We observed differential DNA methylation of the *TXNIP* enhancer between MCI and AD and in the *TXNIP* gene body (2 DMPs). The thioredoxin interacting protein (*TXNIP*) gene product is an endogenous regulator of redox/glucose induced stress and inflammation that is upregulated in the cortex of AD persons.[40] TXNIP co-localizes proximate to Aβ plaques and p-tau neurofibrillary tangles, both hallmarks of AD. The presence of over 50 differentially methylated enhancers that interact with the promoters of genes that are differentially expressed in AD denotes a shared molecular process for DNA methylation changes in enhancers located greater than 5kb from a gene boundary and their putative promoters (*i.e*., >40% of DMPs). To resolve the interplay between cell-type–specific DNA methylation, long-range enhancer-mediated gene expression, and AD pathogenesis future investigations may include single-cell transcriptomics including simultaneous single-cell methylome and transcriptome sequencing (scMT-seq)[56] to measure differential DNA methylation of gene bodies, enhancers, and promoters together with RNA sequencing in serial samples along trajectories of change in cognitive status. DMPs that are shared and those that are unique between independent cohorts of participants with MCI and AD may provide candidate biomarkers for MCI and AD onset and progression in easily accessible blood samples. In particular, machine learning algorithms hold the potential to identify candidate DMPs that distinguish participants by cognitive status and supports testing in longitudinal samples from participants with and without dementia over time.

Our study has multiple strengths. The clinical diagnosis of MCI, AD, and CU phenotypes in the WRAP and WADRC cohorts is based on stringent, well-validated, and consensual clinical and psychometric criteria. WGMS performed with enzyme conversion of unmethylated cytosines rather than harsher sodium bisulfite treatment[13] generated exceptional genomic coverage of 53x (Figure S1).[57] WGMS supports comprehensive resolution of all possible CpGs in the human genome using sequencing by synthesis compared to partial coverage of hybridization-based microarray methods with loci and probes selected for commercial utility.[58] Our sample sizes are substantial and our adjustments for known confounders and corrections for multiple comparisons are robust.

Our study also has limitations. Parallel investigations of differential DNA methylation in persons with MCI and AD from diverse ethnicities is a top priority for future studies. While both MCI and AD have a differential incidence by sex, the statistical model we employed incorporates DNA methylation data from both sexes. All WGMS data herein was generated from bulk populations of white blood cells. Future studies may identify contributions from specific cell types using single cell techniques, such as fluorescence-activated sorting of fixed nuclei. Although our data show that 10% of coding genes are differentially methylated in persons with MCI, the present data alone cannot resolve changes that are causal from those that are compensatory or consequential.

## 5 CONCLUSIONS

The present data provide a substrate for panels of validated methylome biomarkers of MCI and AD in blood configured for translation to clinical and research applications using targeted, rapid, minimally invasive, and cost efficient DMP detection platforms. In particular, DNA methylation biomarkers are cell-based and potentially sensitive to the heterogeneity of syndromes of accelerated changes in cognitive status that may complement plasma biomarker testing. In turn, future genomic, transcriptomic, proteomic, and metabolomic investigations integrated with and by DNA methylation data hold promise to disclose etiologic contributors and to discern personalized AD trajectories at the highest levels of resolution.[59]

## ACKNOWLEDGEMENTS

The authors would like to thank the participants and study personnel who made this work possible.

## CONSENT STATEMENT

The experimental protocol was approved by the Institutional Review Board (IRB) of the University of Wisconsin School of Medicine and Public Health, Madison, WI. All participants signed an IRB-approved informed consent.

