## Supplementary material for "Whole genome methylation sequencing in blood from persons with mild cognitive impairment and dementia due to Alzheimer’s disease identifies cognitive status": Figure S1

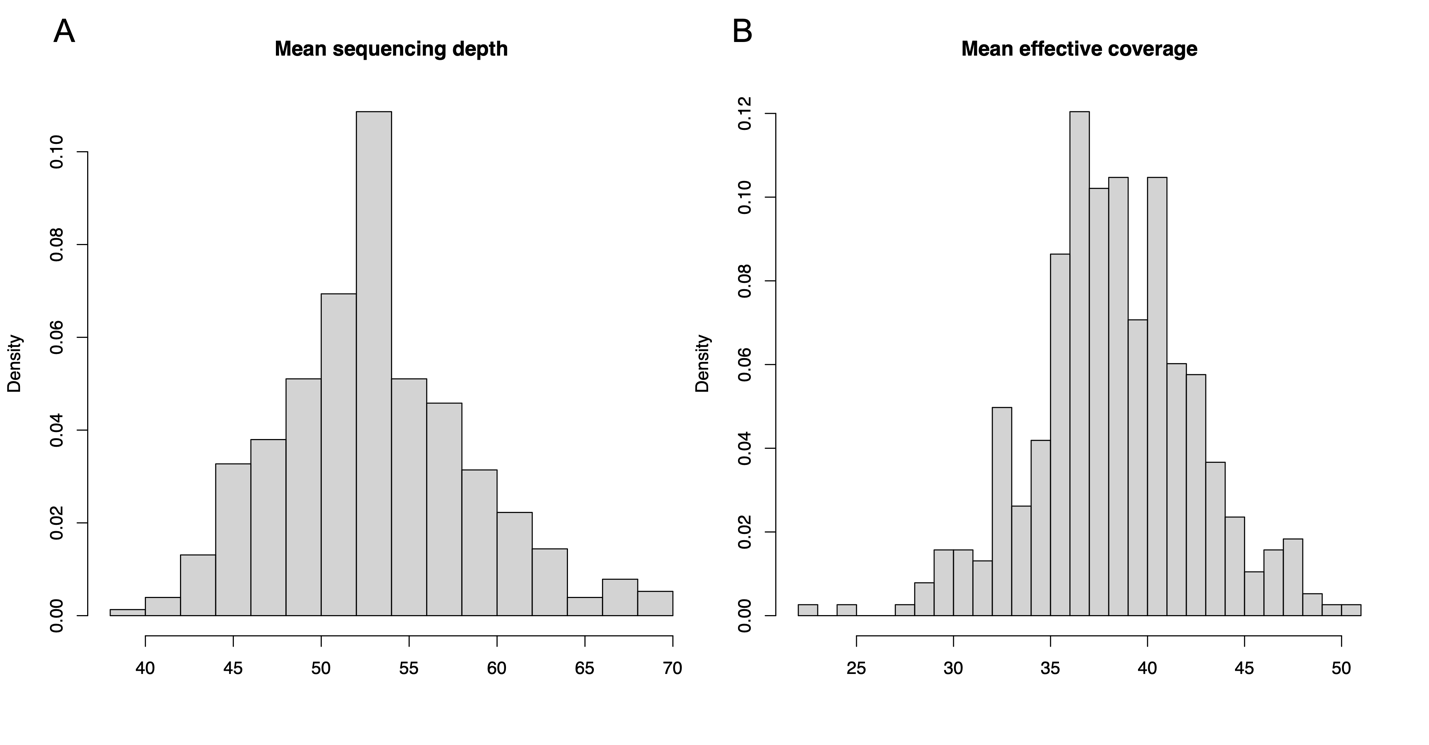

**Figure S1:** Distribution of average sequencing read depth and effective coverage.

(A) The average sequencing depth per participant sample (53x) is defined as the sequence read length multiplied by the number of sequence reads divided by the haploid genome length after data processing[1]. (B) The effective coverage of 38x is defined as the average number of sequence reads over a specific CpG after merging across strands. Density (y-axis) indicates the relative amount of genomic sequence read depth and effective coverage (*i.e.* the number of informative reads for methylation after merging across strands, respectively). Sequencing depth and effective coverage are not identical because not all CpGs have measurements on both strands after data processing.
