## Supplementary material for "Whole genome methylation sequencing in blood from persons with mild cognitive impairment and dementia due to Alzheimer’s disease identifies cognitive status": Figure S2

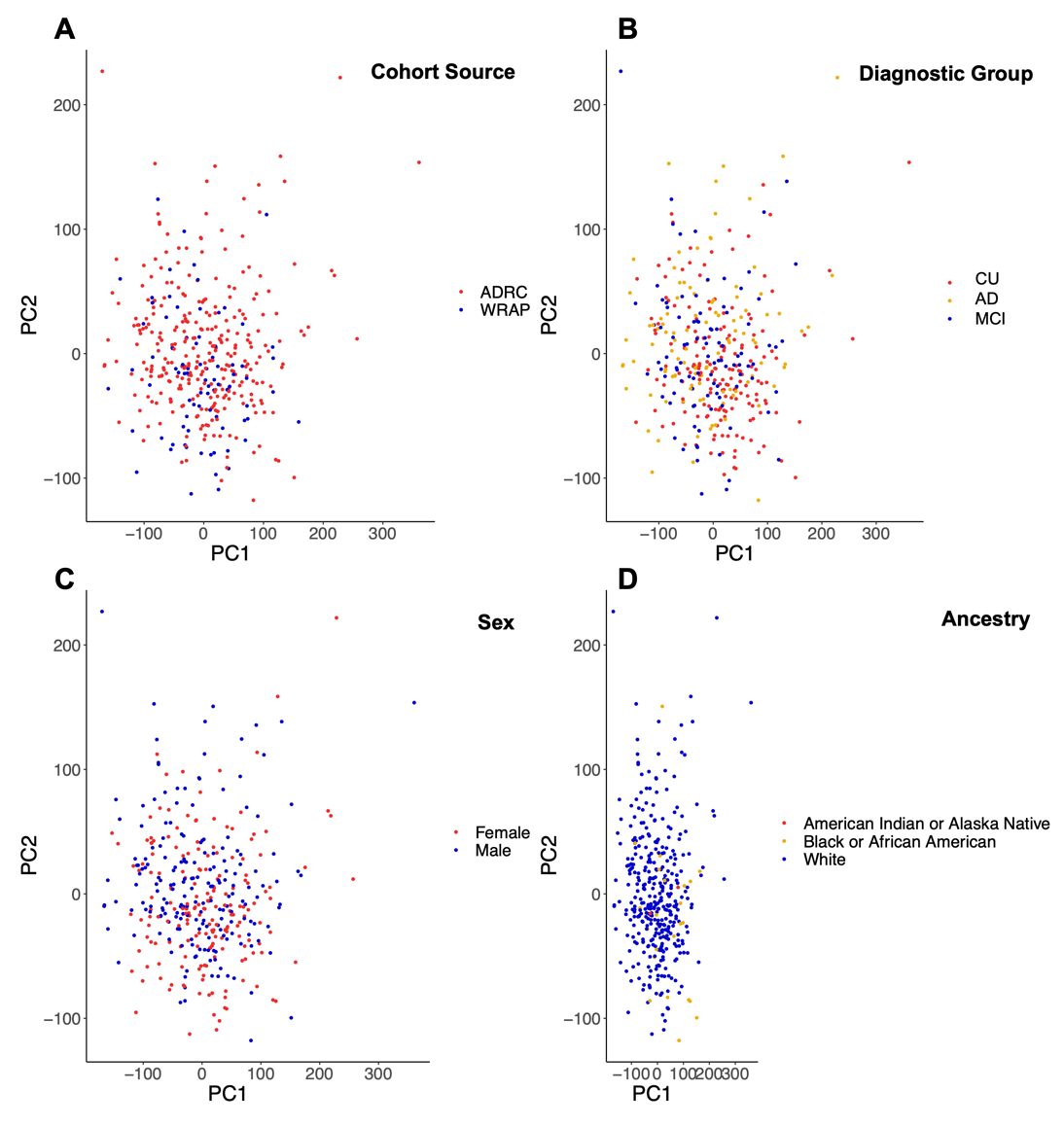

**Figure S2:** The first 2 principal components (PC) of the 5% most variable CpGs on chromosome 1. PC-1 on the x-axis accounts for 5.1% of total variance and PC-2 on the y-axis accounts for 2.5% of total variance. Plot represent PC outputs from cohort source (A, WADRC and WRAP), diagnostic group (B, AD, MCI, or CU), sex (C), and self-reported ancestry (D). Lack of clustering for data cohort indicates the absence of batch effects. The lack of clustering for diagnostic status, sex, and self-reported ancestry indicates that PCs alone do not capture variability due to diagnostic status, sex, or ancestry.
